# *Denovo* Identification of Cross-linked Peptides via Isotope Coded Linkage Tags

**DOI:** 10.1101/2020.04.12.037614

**Authors:** Harsha P. Gunawardena

## Abstract

An isotope labeled cross-linker (asymmetric d4-DTSSP) was developed to streamline the efforts required for the detection of cross-linked peptides. The cross-linking and mass spectrometry strategy we call Isotope Tagging of Interacting Proteins (iTIP) has improved the specificity of detecting cross-linked peptides and the accurate identification of the interacting peptide sequences via the incorporation of isotopic signatures that are readily observed in the MS/MS spectra. All tryptic peptides derived from the cross-linking reactions of a protein complex are first subjected to ETD-MS^2^ which results in the facile cleavage of the cross-linker at the disulfide bond and the release of inter-linked polypeptide chains that are detected as a pair of peaks (doublets) in the MS^2^ spectrum. The constituent peptide halves that are tagged by the heavy/light ends of the cross-linker are easily mass-selected from all other fragment ions, and each polypeptide half is then subjected to CID or HCD-MS^3^ for identification. The MS^3^ spectra are subjected to conventional database search strategies available for the sequencing of linear or non-cross-linked peptides. The confident identification of each polypeptide is further assisted by the presence of a stable isotope labeled fragment ions that localizes the cross-linked site on the polypeptide sequence.

## INTRODUCTION

A majority of gas-phase dissociation reactions of protonated cross-linkers lack the sensitivity and specificity required to fragment the cross-linker at a desired location while giving rise to diagnostic fragment ions; only a few reports have attempted to address the issue of selectivity^1-8^. Reid and co-workers described a cross-linker containing a sulfonium ion that showed specific fragmentation of the C-S bond during CID.^9^ Brodbelt and co-workers reported the preferential cleavage of N-N hydrazone bond of a bis hydrazone cross-linked peptides by ETD.^10^ In both reports, despite the fact that dominant fragment channels arose from a specific cross-linker cleavage, the inability to readily distinguish fragment ions unique to the cross-linker from backbone fragment ions makes cross-linked peptide diagnosis challenging. Bouchers and co-workers have reported an isotope coded CID cleavable cross-linker.^7^ Although their cross-linker has diagnostic fragment ions, the biotin moiety introduced to their cross-linker adds substantial mass to the resulting interlinked peptides. Here, we have synthesized “asymmetric d4-DTSSP” a cross-linker with a relatively smaller mass that can be selectively fragmented at a specific location, while allowing the dissociated cross-linked fragment ions to be distinguished easily from other backbone fragments. The stable isotopes are incorporated in DTSSP via asymmetric positioning of four deuterium labels on two of the four methylene carbons, giving rise to a mass asymmetry across the two-fold axis of the S-S bond. Further, d4-DTSSP is structurally analogous to the commercially available DTSSP reported elsewhere.^11^ However, the unique asymmetric isotope tagging signature of our cross-linker can be detected via the selective electron transfer dissociation of the disulfide bond^12^ of the cross-linker. The asymmetric nature of the cross-linker can render a doublet of reporter ions with a nominal mass off-set of 4 Da per unit charge for each inter-linked peptide half, due to the equal likelihood of orienting the light and heavy ends of the cross-linker during the reaction. The h4 and d4 doublets (light and heavy-isotope tags attached to the constituent peptides) are therefore specific for only cleavage products associated with the disulfide cross-linker and hence diagnostic of each interacting peptide pair. The small mass off-set makes these fragment ions predictable, easy to visualize, and easily distinguishable from other ETD product ions^13^ such as peptide backbone products, side-chain losses, and charge reduced species which are single peaks. Each of the peptides cleaved at the S-S bond is structurally interrogated via MS^3^-CID where the product ion spectra of each peptide half forms the basis for their identification and hence the identification of an interacting protein. We have named this new cross-linking strategy that identifies interacting proteins as Isotope Tagging of Interacting Proteins (iTIP).

## EXPERIMENTAL SECTION

### Materials

All chemicals and solvents were of the highest grade and used without further purification. 3-Mercaptopropionic acid, 2,2’-dithiodipyridine and dimethylformamide (DMF) were purchased from Acros. Dicyclohexylcarbodimide (DCC) and ethanol were purchased from Sigma-Aldrich. *N*-Hydroxysulfosuccinimide sodium salt (NHSS) was purchased from Fluka. Acetic acid and ethyl acetate were purchased from Fisher Scientific. 2,2’,3,3’-Tetradeuterium-3-mercaptopropanoic acid was purchased from Creative Molecules Inc.

### Synthesis of Asymmetric d4-DTSSP

Figure 1 shows the overall synthetic scheme for generating asymmetric d4-DTSSP. The disulfide **1** was synthesized as described by Xie et al.^14^ Disulfide **1** (16.9 mg, 79 μmol), THF (0.4 mL), H_2_O (0.3 mL) and 2,2’,3,3’-Tetradeuterium-3-mercaptopropanoic acid (83 μL, 1 M in D_2_O, 83 μmol) were added successively to a vial at room temperature. The reaction mixture was stirred for 1 hr and then concentrated under reduced pressure. Flash chromatography (CH_2_Cl_2_ : MeOH = 95 : 5 with 0.4 % formic acid) over silica gel gave compound **2** as a white solid (9.3 mg, 55 %). NHSS (8.7 mg, 40 μmol) and DCC (8.3 mg, 40 μmol) were added successively to a solution of compound **2** (4.3 mg, 20 μmol) in DMF (0.3 mL) at room temperature. The resulting solution was stirred for 4 hrs and then subjected to centrifugation. The supernatant was collected and 1.2 mL of ethyl acetate was added. The resulting mixture was centrifuged. The precipitate was collected and washed by ethyl acetate three times. After drying by SpeedVac, the cross-linker **3** was obtained as a pale white solid (7.0 mg, 57 %). Figure 1a describes the synthetic scheme used to generate d4-DTSSP. The synthetic product was fully characterized by NMR ^1^H NMR (CDCl_3_, 400 MHz) δ 3.11-3.15 (m, 2H), 3.19-3.26 (m, 4H), 3.40 (d, *J =* 9.2 Hz, 1H), 3.44 (d, *J =* 9.6 Hz, 1H), 4.33 (d, *J =* 8.8 Hz, 1H), 4.52 (d, *J =* 8.8 Hz, 1H), and high resolution mass spectrometry data (Figure 1b) that compares commercially available DTSSP, d8-DTSSP and newly synthesized asymmetric d4-DTSSP.

**Figure 1.**
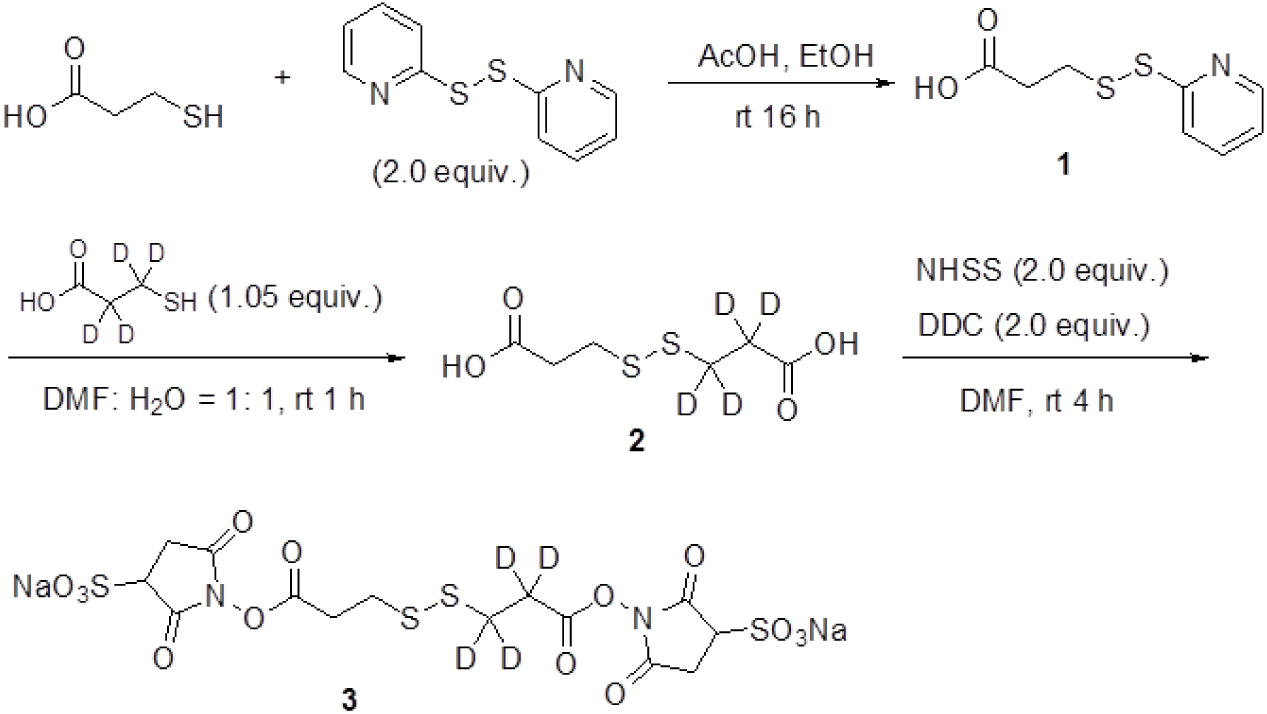
The synthetic procedure for generating d4-DTSSP.

### Protein Cross-Linking and Proteolysis

Protein cross-linking was carried out on ubiquitin and holo myoglobin using the manufacturer’s protocol described for DTSSP cross-linkers (Pierce Inc.) The protein solution was buffer exchanged to 25 mM ammonium bicarbonate (ABC) and trypsinized at an enzyme to protein mole ratio of 1:50 at 37 °C in a 3M spin-filter device (Millipore Inc.,). Peptides were then desalted using PepClean spin filters (Pierce Inc.) and reconstituted in an aqueous solution of 0.1% formic acid.

### Mass Spectrometry Analysis

Data dependent LC-MS/MS was carried out on a LTQ-Orbitrap Velos mass spectrometer (Thermofisher Scientific) coupled to an Eksigent 2D nano ultra LC system as described previously.^15^ The targeted experiments consisted of ETD-MS/MS, followed by a minimum of two CID-MS/MS scans or two HCD-MS/MS scans for each polypeptide light/heavy chain. Mass spectra were processed by filtering MS^3^-CID or -HCD spectra, and peptide identification was performed on the filtered MS^3^ spectra by Mascot (Matrix Science Inc.) against a human Uniprot database (V3.63). Peptides were confidently identified using a target-decoy approach^16, 17^ at a 1% false discovery rate (FDR). A precursor ion mass tolerance of 10 p.p.m. and a product ion mass tolerance 0.5 Da were used during the initial search with a maximum of two missed trypsin cleavages. Variable modifications included methionine oxidation and customized cross-linker associated modifications: d4-S, d4-SH, d4-S, d4-SH at lysine and N-terminus. All search results were filtered for precursor masses to be within a 6 p.p.m mass accuracy.

## RESULTS AND DISCUSSION

### Rationale for Cross-linker Design and Isotope Tagging of Interacting Proteins (iTIP) Strategy

It has been demonstrated that electron transfer dissociation (ETD) provides facile cleavage of peptide backbone bonds^13, 18^, disulfide bonds of oxidized cysteine residues^,12^ and a number of cross-linked polypeptides.^10^ Based on these observations, we rationalized that an ETD cleavable cross-linker consisting of a disulfide bond could be quite useful in studying inter-linked polypeptides in cross-linking-based tandem mass spectrometry experiments. Commercially available disulfide containing cross-linkers such as DTSSP and DSP, and their deuterated analogs are structurally symmetrical across the disulfide bond: with a two-fold symmetry, each constituent peptide chain results in identical mass-tags in ETD-MS^2^ spectra. Figure 2a shows the structure of our asymmetric d4-DTSSP cross-linker with a design strategy based on a mass off-set or (Δm=4 Da) around the S-S bond. The bidirectional orientation of the cross-linker facilitates two fragments in the ETD spectrum for each constituent chain by tagging of heavy c-b segment or light a-b segment of the cross-linker to an amine group of an N-terminus or lysine residue of a peptide via acylation chemistry. The cleavage site is shown as b or the disulfide bond of the cross-linked peptide. Figure 2b shows the ETD product ion spectrum of an inter-linked polypeptide resulting from a cross-linking reaction of ubiquitin and commercially available DTSSP. The product ions consist mainly of α and β chains that are single isotope clusters resulting from the direct cleavage of the DTSSP disulfide bond. Figure 2c shows the ETD product ion spectrum of the same constituent polypeptide chains inter-linked with asymmetric d4-DTSSP. The product ion spectrum looks quite similar to 1b. However, due to the asymmetry on either side of the S-S bond, and due to the bidirectional orientation of the cross-linkers during the cross-linker reaction step, we observe a doublet of peaks for both α and β chains, labeled as α-L/α-H and β-L and β-H. We rationalize that cross-linkers with asymmetric labeling are analytically useful as they encode isotopic tags of cross-linked product ions to be readily distinguished from other cleavages due to dissociation of the cross-linker. The mass spectra of isotope coded cross-linked peptides consist of two distinct fragment ions for each constituent peptide chain with a specific mass signature that is observed as a doublet of peaks in the ETD-MS^2^ spectrum. Figure 3a shows an ETD-MS/MS of [M+4H]^4+^ precursor ion m/z = 628.3 Da of an interlinked cross-linked polypeptides that was derived from d4-DTSSP cross-linking reactions with holomyglobin. Unlike Figure 1b, where disulfide cleavage products were the dominant peaks, the ETD products associated with the disulfide cleavage products are less conspicuous. The product ions comprise of a distribution of charge reduced species [M+4]^3+•^, [M+4]^2+••^, a series of backbone cleavage product ions that gives rise to c- and •z-type ions, and three doublets of fragment ions corresponding to the cleavage of isotope coded disulfide-linked cross-linker. The doublet peaks are the only diagnostic features in the mass spectrum that is informative of an interlinked polypeptide with two chains. The doublet of peaks of m/z = 759.4, and m/z = 763.4 have a nominal mass difference of 4 Da, corresponding to the ΔM generated by the dissociation across the disulfide of an asymmetric d4-DTSSP cross-linker. Close examination of the isotope clusters of both these peaks shows an isotope spacing of ∼ 1 Da indicating that they are singly charged fragment ions. Similarly, the doublet of m/z = 875.2 and m/z = 877.2 are separated by a nominal mass difference of 2 Da with each peak having an isotope spacing of 0.5 Da. The isotope spacing indicates that a doubly charged fragment has a ΔM of 4 Da generated via a cleavage across a d4-DTSSP cross-linker. These diagnostic doublets of peaks are therefore directly related to a peptide cross-linked by an asymmetric d4-DTSSP. Further, both these doublets we believe are complementary fragments generated via the single dissociation of charged reduced radical cations: [M+4]^3+•^ at the disulfide bond since cleavage resulted in the conservation of the overall charge state of the charged reduced radical cation. The constituent peptide chains with their corresponding charge states for doublet pairs are labeled as [α-L]^+^/[α-H]^+^ and [β-L]^2+^/[β-H]^2+^. Further, the doublet of m/z = 1750.7 and m/z = 1754.7 has a nominal mass difference of 4 Da with each peak having an isotope spacing of 1 Da, again indicating a singly charged fragment ion corresponding to the isotope coded product ions of the cross-linker. We believe the generation of these doublet peaks [β-L]^+^/[β-H]^+^ was due to the sequential charge reduction of the doubly charged doublets [β-L]^2+^/[β-H]^2+^. The iTIP approach readily generated diagnostic fragment ions from cross-linked peptides that can be structurally interrogated via a CID step that is more suitable for doubly and singly charged peptides halves constituting the cross-linked peptide halves.

**Figure 2.**
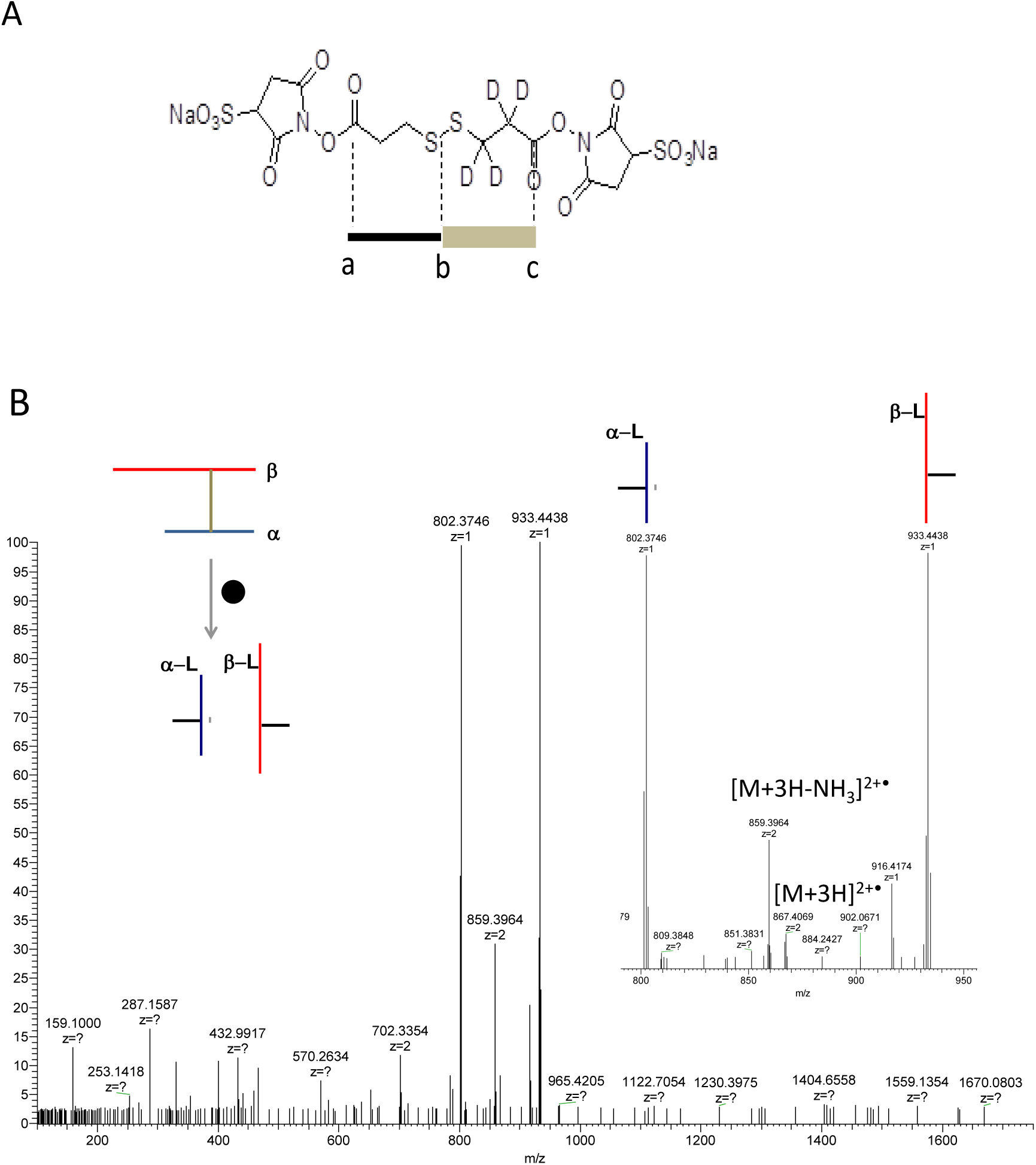
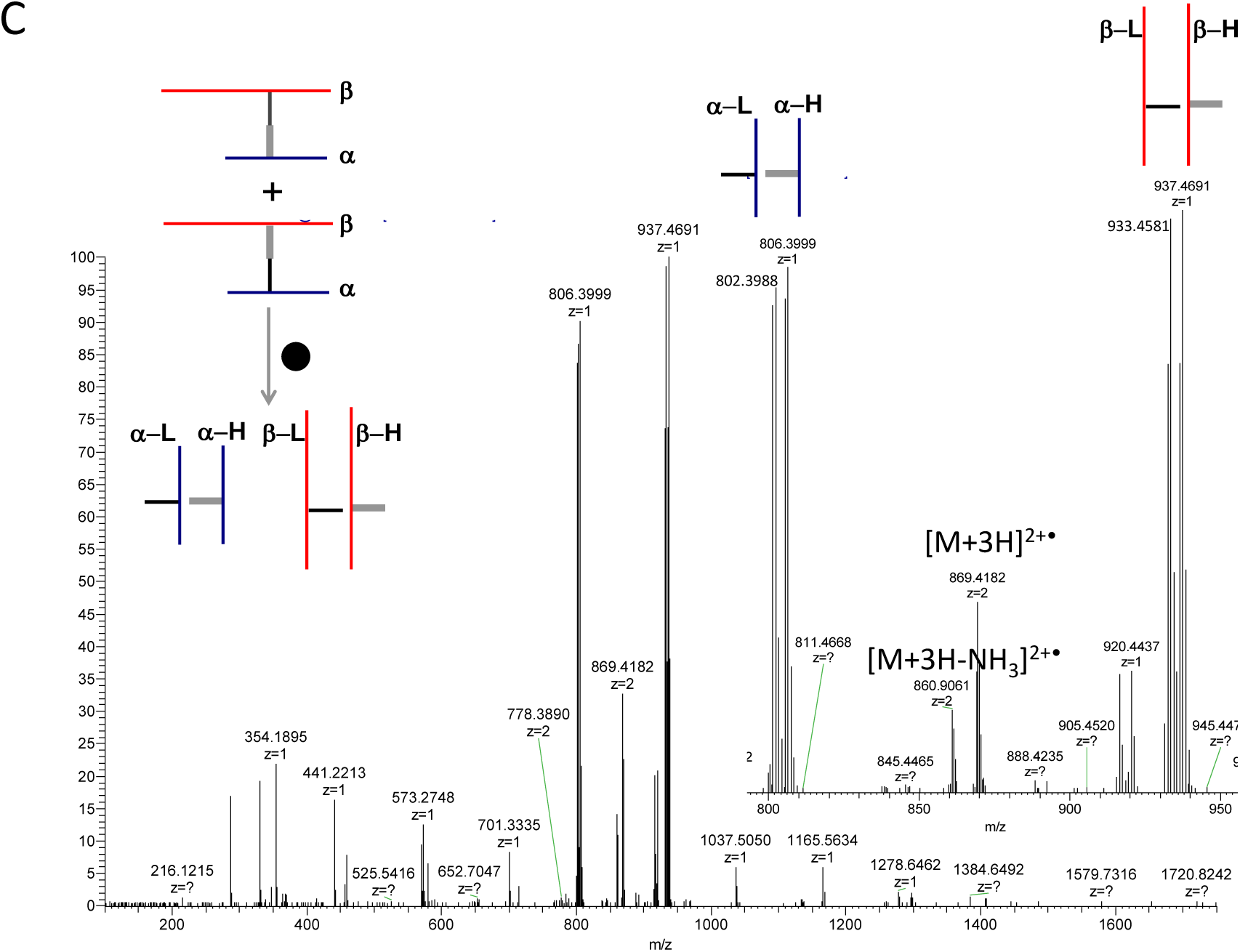
(A) Asymmetric d4-DTSSP structure showing amine reactive sites a, c and ETD cleavable site b. (B) ETD-MS/MS spectrum of inter-linked peptide between the reaction of ubiquitin and DTSSP. Insets shows dissociation of the cross-linker to yield the constituent α and β peptides (C) ETD-MS/MS spectrum of the same polypeptide pairs intelinked via asymmetric d4-DTSSP. Insets shows dissociation of the cross-linker to yield the constituent α-L and α-H polypeptide and β-L and β-H polypeptide

**Figure 3.**
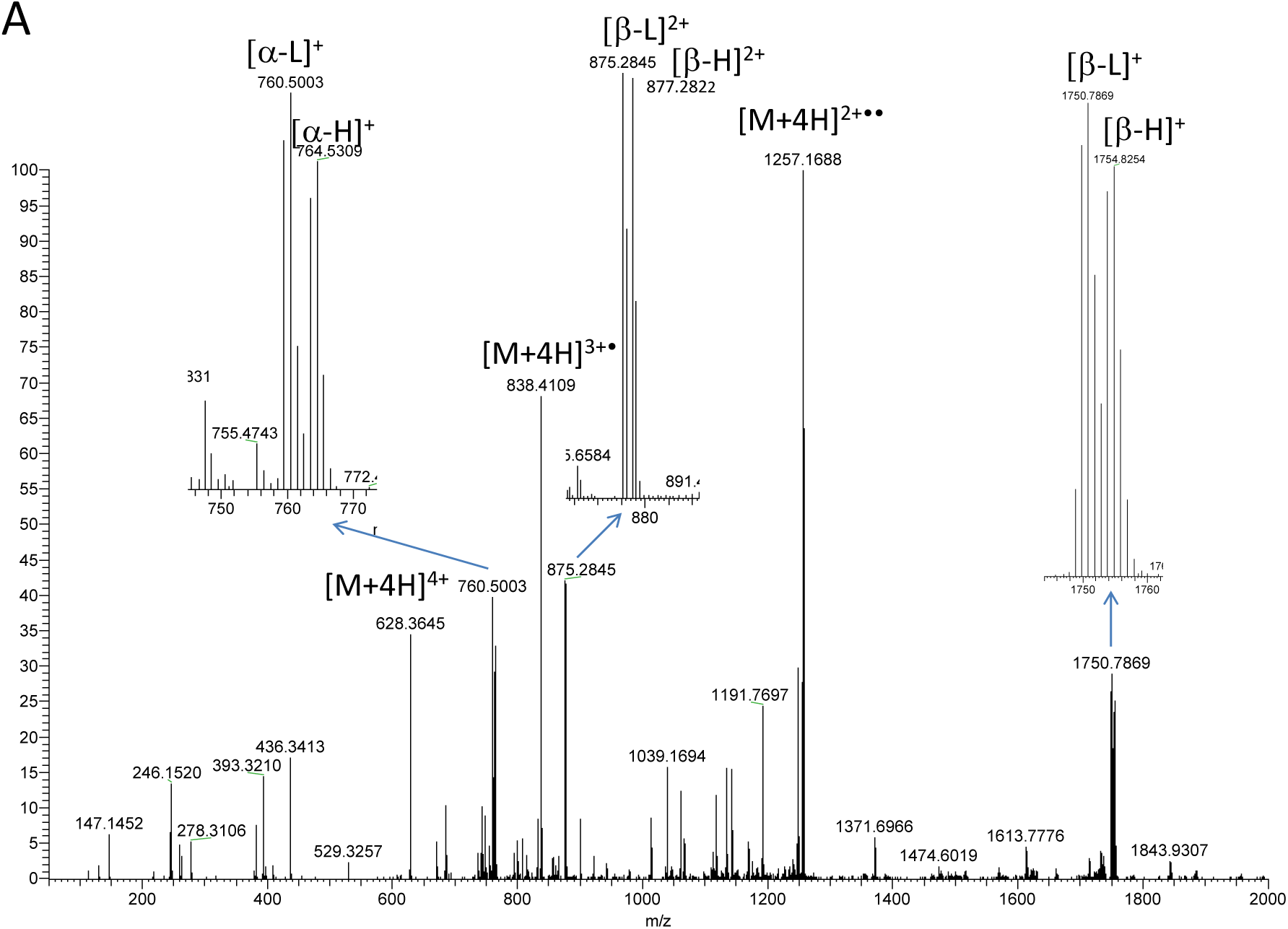
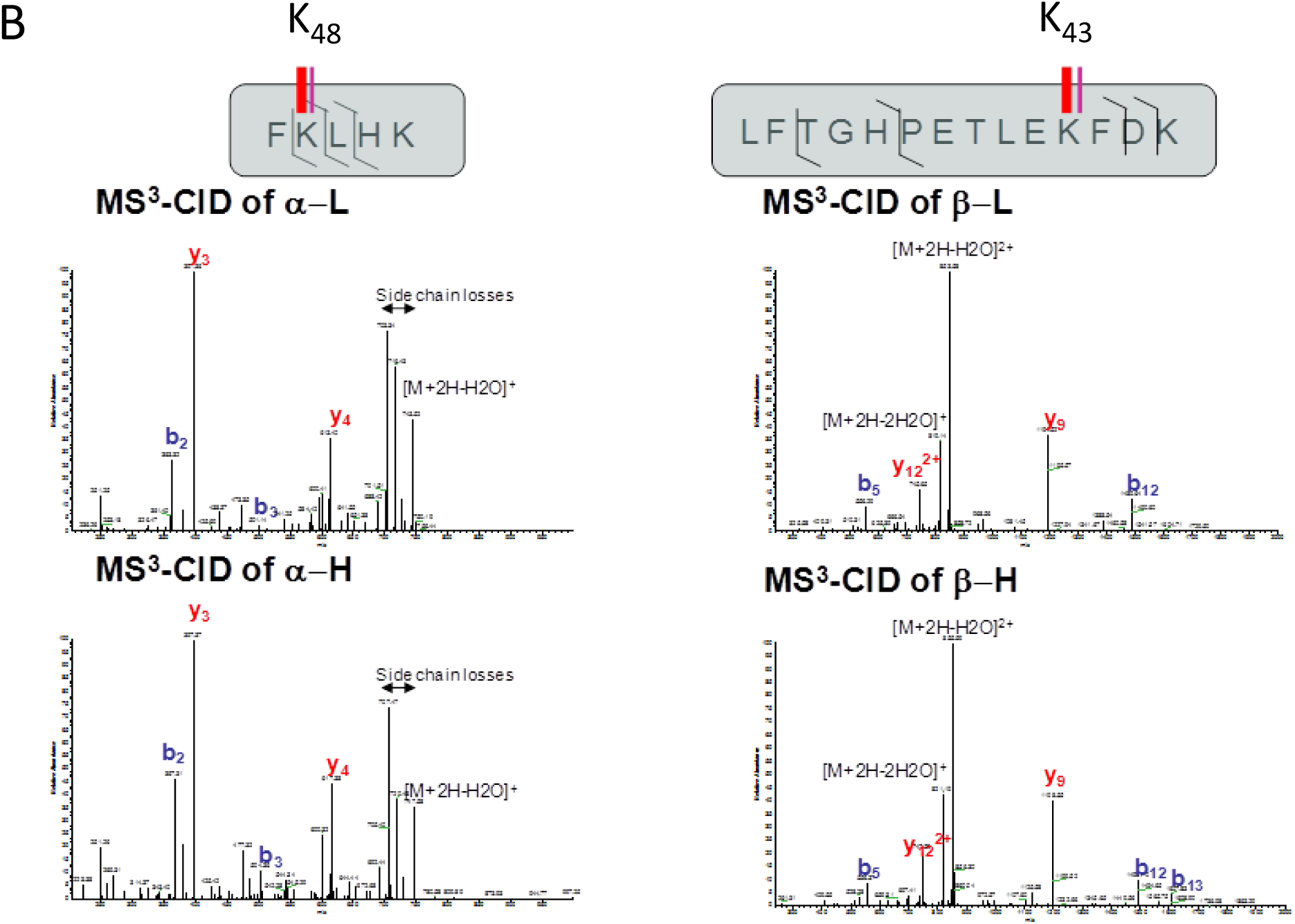
(A) ETD-MS/MS spectrum of a quadruply charged inter-linked peptide resulting from the cross-linking reaction between holomyoglobin and asymmetric d4-DTSSP. Insets shows show the dissociation products associated with the cross-linker that is seen as an isotope coded mass tag with each constituent chain labeled as α-L and α-H doublet and β-L and β-H doublet (B) CID-MS3 of α-L, α-H, β-L, and β-H and the corresponding sequence map of each polypeptide depicting CID fragmentation and location of lysine residues that are cross-linked

### Identification of Interaction Peptide Pairs

The selective identification of cross-linked peptides becomes straightforward and streamlined by the MS^3^ step. Here we show that the peptide doublets generated by ETD can be sequenced by conventional database assisted peptide search strategies. For example, the doublet peaks that are shown in Figure 3a were structurally interrogated via CID-MS^3^ and the spectra were subjected to a database assisted sequence determination of the interlinked polypeptides. The iTIP strategy also allows MS^3^-CID of both heavy/light chains of each peptide sequence which provides an additional degree of confidence in their identification as interacting proteins or interaction sites within a protein. Figure 3b shows the resulting CID product ion spectra for the α-chain_L,_ α-chain_H_, β-chain_L_, and β-chain_H_ of the constituent cross-linked peptide chains of the two interacting lysine residues of holomyglobin. Each type of chain is discernable by its respective isotope spacing encoded in Light-H4 methylene and Heavy-d4 methylene. The sequence for each constituent polypeptide ion was identified using a standard database search with dynamic lysine modification as described in the experimental section. The polypeptide sequences were identified as myoglobin and the cross-linked tags were localized to K43 and K48. The constituent peptide sequences are referenced back to the corresponding MS^2^-ETD spectrum to make a pair-wise interaction map that shows that K43 and K48 of myglobin was cross-linked by d4-DTSSP and the interaction was within a distance of 12 Å or less.

## CONCLUSIONS

We call this overall strategy of identifying and localizing putative lysine residues that belong to a single protein or complexes as Isotope Tagging of Interacting Proteins (iTIP). The iTIP procedure starts with cross-linking of whole protein molecules or complexes with d4-DTSSP, followed by proteolysis and tandem mass spectrometry involving ETD of the inter-peptide cross-links to generate isotope tagging patterns that serve as structural surrogates for identifying protein-protein interactions. The iTIP technology is amenable to identifying lysine residues that constitute the interacting surfaces of proteins with a high degree of confidence as demonstrated by CID-MS/MS of both light and heavy forms of the same peptide sequence. We envision that iTIP could be used in many structural biology and structural proteomics applications where asymmetric d4-DTSSP analogs of different linker lengths can be leveraged to streamline cross-linking-based mass spectrometry experiments.

## ACKNOWLEDGEMENT

The authors would like to thank the University Cancer Research Fund in support of this research. HPG conceived the idea, designed and performed all MS experiments and wrote the paper. The cross-linker and methodology reported is patented under US patent US 9,562,010 B2 and assigned to the University of North Carolina at Chapel Hill.

---

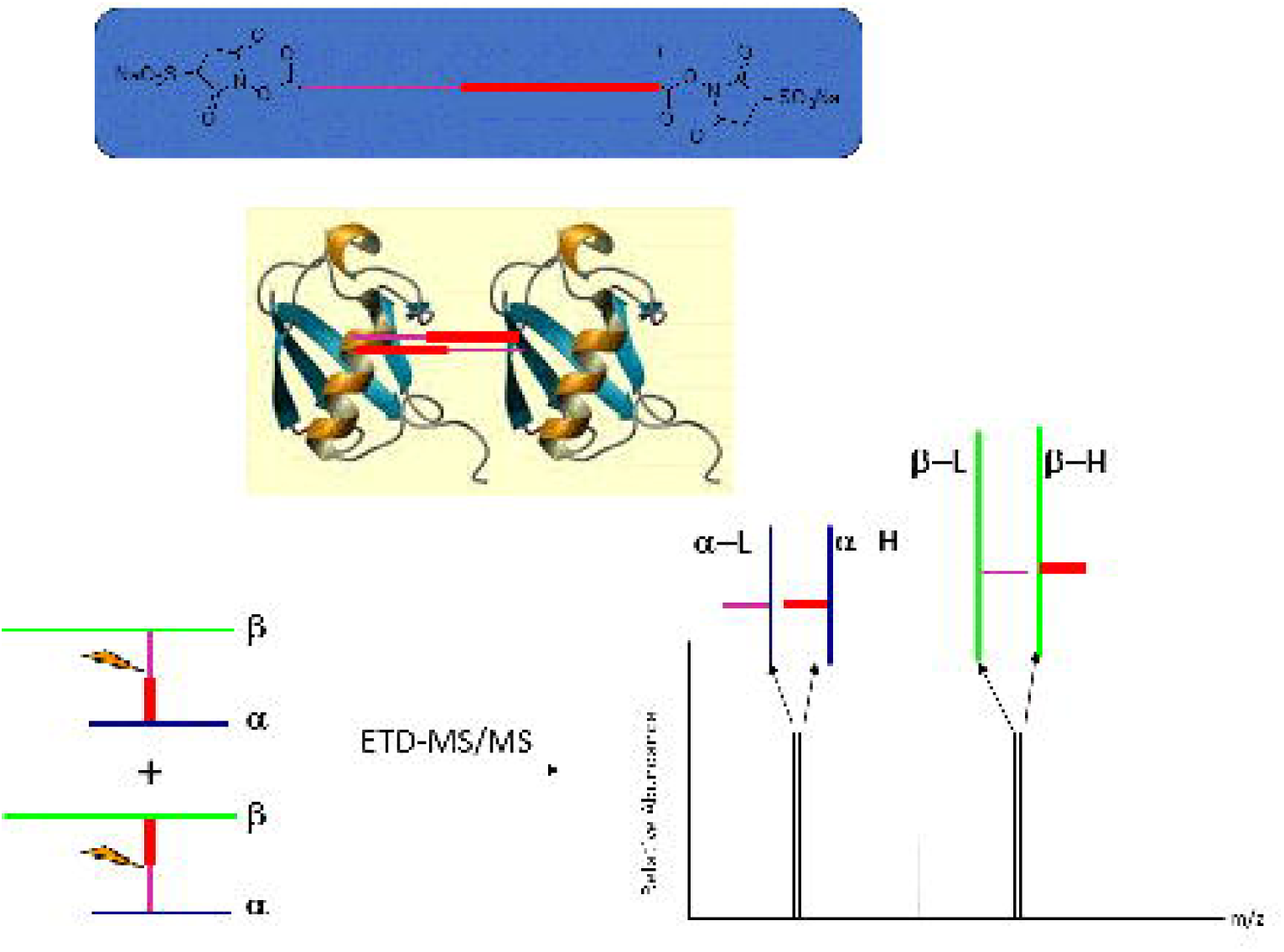

